# A thioester-thiol exchange reaction in *Salmonella* Invasion acyl carrier protein transfers the 4’PP-bound acyl chain to a distal cysteine

**DOI:** 10.64898/2026.10.23.761286

**Authors:** Rashima Prem, Garima Garima, Debodyuti Sadhukhan, Monica Sundd

## Abstract

*Salmonella* Typhimurium encodes two functionally distinct acyl carrier proteins: AcpP, involved in fatty acid synthesis, and the invasion acyl carrier protein (IacP), which mediates the acylation of SipB (*Salmonella* invasion protein B), a modification essential for SipB maturation and efficient host invasion. Despite growing insights, the IacP-dependent machinery governing acyl chain transfer remains largely enigmatic. Therefore, a multidisciplinary approach combining biochemistry and NMR was employed. A D35NIacP mutant (IacP^N^) that expresses in the apo-form was enzymatically converted into acyl-IacP^N^ intermediates. Unexpectedly, the ^1^H^15^N HSQC spectra of the acyl-IacP^N^ intermediates displayed two sets of resonances for several of the amides. The first was a less intense set of peaks, positioned close to the holo-peak, displaying a progressive chemical shift change with increasing acyl-chain length, reminiscent of the 4’-phosphopantetheine (4’-PP)-bound conformer of type II ACPs. The second, a more intense, acyl-chain-length-independent, previously uncharacterized set of peaks, was clustered near the longer 4’PP-bound acyl-chain conformers (C14-). The origin of this new conformer was investigated by generating three cysteine-to-alanine mutants, C16A, C54A, and C69AIacP^N^. The C54AIacP^N^ mutant displayed a single set of peaks in the acyl-IacP^N^ intermediates, corresponding to the 4’PP-bound IacP^N^ conformer, establishing the critical role of Cys 54 in generating the uncharacterized peak and identifying it as a Cys-bound conformer. The acyl-S-carboxyamidomethyl-IacP^N^ intermediates also displayed a single conformer, supporting the role of a cysteine in giving rise to the new peak. Collectively, our studies uncover a unique thioester-thiol exchange reaction in which the 4’PP-tethered acyl chain is chemically transferred to a distal cysteine (Cys 54). Given that Cys 54 lies within the SipB interaction interface of IacP, this acyl chain relay process may contribute to IacP-mediated SipB acylation and *Salmonella* virulence.

## Introduction

*Salmonella* Typhimurium is a facultative intracellular pathogen and a leading cause of gastroenteritis in humans. More recently, persistent *Salmonella* infection has been linked to gallbladder cancer and bloodstream infections, underscoring the need to elucidate its underlying mechanism of pathogenesis (*1–3*). The bacterium invades the intestinal epithelial cells using a type III secretion system (T3SS) encoded by the *Salmonella* pathogenicity island-1 (SPI-1). Notably, the *iacp* gene, which encodes the invasion acyl carrier protein (IacP), is located within the SPI-1 locus and is cotranscribed with the *sipBCDA* operon, which encodes key T3SS components (*4*). Functional studies have demonstrated that the deletion of the *iacp* gene significantly impairs *Salmonella* entry into the intestinal epithelial cells, and hence the name ‘invasion acyl carrier protein’ or IacP (*5*). Mechanistically, IacP is required for the acylation of SipB (*Salmonella* invasion protein B), a posttranslational modification essential for SipB maturation and *Salmonella* virulence.

In addition to the atypical IacP, *Salmonella* Typhimurium also encodes a canonical AcpP, which functions in the type II fatty acid synthesis pathway (FAS). The *Salmonella* AcpP sequence is identical to that of *E. coli* AcpP, and consequently, the two proteins share the same structure. Besides these two pathogens, canonical type II ACPs have been reported in most prokaryotes, including *M. tuberculosis, Plasmodium falciparum, Helicobacter pylori, as well* as a few eukaryotic organelles, such as plastids and mitochondria (*6–9*). The fundamental difference between Type I & type II ACPs lies in the organization of their biosynthetic machinery and the mechanism of acyl chain sequestration. In the type I pathway, various components/enzymes exist as domains of a single, large, multifunctional, multidomain complex. This type of ACP is present in *Mycobacterium tuberculosis*, *Saccharomyces cerevisiae*, *Candida albicans*, and *Homo sapiens* (*10–12*). In addition, several polyketide synthases and ribosomal peptide synthases also employ the type I pathway to synthesize diverse biomolecules (*13, 14*). In contrast, the type II pathway, often referred to as the dissociated pathway, consists of discrete, independently functioning enzymes. Within this pathway, the central player-the acyl carrier protein (ACP) exhibits unique conformational plasticity to shield the growing acyl chain from premature hydrolysis and transfer into the active-site cavity of the pathway enzymes (*15*). Thus, most canonical type II ACPs sequester the lengthening acyl chain within their hydrophobic cavity, releasing it into the active site of their pathway enzymes *via* a unique switchblade mechanism. In contrast, type I ACPs lack the chain-flipping machinery (*16*). Instead, the acyl chain tethered to the type I ACP remains associated with the hydrophobic residues present within the crevices and grooves on the ACP surface to prevent thioester bond hydrolysis (*17*). Together, the two ACPs play indispensable roles in several biosynthetic pathways, including fatty acid synthesis (FAS), polyketide synthesis (PKS), and nonribosomal peptide synthesis (*18–20*).

Across all genera, a type I or a type II machinery is conserved, although a subset of organisms, including *M. tuberculosis, Homo sapiens,* and most mammals, possess both. In *M. tuberculosis*, the type I and type II pathways function in a coordinated manner. The type I pathway synthesizes a C12-C16 acyl chain precursor, which is subsequently elongated to C24-26 by the type II pathway, ultimately producing complex mycolic acids (*21*). In mammals, however, two independent pathways coexist in different cellular compartments: a type I FAS in the cytosol, which synthesizes 90% of cellular fatty acids, and a type II pathway in the mitochondria (*22*). The mitochondrial pathway synthesizes long-chain fatty acids for mitochondrial membrane biogenesis and C8 fatty acids, which serve as precursors for lipoic acid synthesis (*9*). Beyond their biosynthetic role, mitochondrial ACPs have emerged as important structural components of large multiprotein complexes, including mitoribosomes, Fe-S cluster biogenesis complexes, and electron transport chain complexes I, II, and III, which are essential for mitochondrial metabolism and function (*23, 24*).

Despite low sequence identity, most ACPs (types I and II) adopt a conserved four-helical bundle architecture (*17*). They are expressed as inactive apo-protein and subsequently converted to their functional holo-form through posttranslational modification. This activation involves the transfer of a 4’-phosphopantetheine group from coenzyme A to a conserved serine (Ser 36) by the catalytic action of a 4’-phosphopantetheinyl transferase (PPTase). The modified serine is located within a ‘DSL’ motif at the beginning of helix II, critical for PPTase recognition and phosphopantetheinylation. The holo-ACP so formed enters the biosynthetic cycle, where fatty acyl-, peptide-, and polyketide intermediates are assembled on the free thiol of its phosphopantetheine (4’-PP) arm. This prosthetic group acts as a flexible swinging arm that shuttles the pathway intermediates between the active sites of the partner enzymes. During fatty acid synthesis, the growing hydrocarbon cargo undergoes multiple cycles of condensation, reduction, dehydration, and a second reduction step, each catalyzed by distinct enzymes of the fatty acid synthesis pathway. These iterative cycles continue until the acyl chain reaches a typical length of C16-C18. At this stage, the mature acyl chain is recognized and cleaved by the catalytic action of an acyl-ACP thioesterase, releasing the fatty acid (product) to perform its destined cellular function(*25*). Throughout the elongation cycle, ACP dynamically interacts with the pathway enzymes primarily through its helix II, also referred to as the ‘universal recognition motif’, a protein-protein interaction surface conserved in all type II FAS ACPs(*26*).

To date, most studies have focused on the canonical FAS ACPs of the type I or type II pathway, whereas the structure, dynamics, and function of noncanonical ACPs remain largely unexplored. Nevertheless, noncanonical ACPs have been identified in a wide range of organisms and play diverse roles. For example, *Enterococcus faecalis* encodes an atypical ACP that scavenges exogenous fatty acids (*27*). *Ralstonia solanacearum* has six ACPs(*28*). *Arabidopsis* has multiple ACP isoforms that are expressed in a tissue-specific manner in response to environmental stress (*29*). Members of the rhizobial group of bacteria, three distinct ACPs coexist, namely NodF, AcpXL, and RkpF to synthesize complex membrane proteins involved in nodulation (*30*). *Streptomyces* species encode eight different ACPs to synthesize a complex range of secondary metabolites (*31*). Gram-negative enteropathogens such as *Salmonella* spp., *Shigella flexneri*, and *Yersinia enterocolitica* likewise possess a specialized invasion acyl carrier protein (IacP) for host invasion and virulence (*32, 33*). Although these specialized ACPs are clearly important for the physiology and virulence of these organisms, they remain poorly characterized, underscoring the need for mechanistic studies on select ACPs. *Salmonella* IacP represents one such atypical ACP that facilitates the maturation of its invasion protein B (SipB) through a mechanism that remains unclear (*34, 35*). To address this gap in our understanding of IacP function, a comprehensive investigation of IacP and its acyl intermediates has been undertaken. Central to our approach was NMR, one of the few techniques capable of providing atomic-resolution insights into the transient, sparsely populated, or otherwise invisible conformational states (*36*). Importantly, a single NMR experiment can capture multiple conformational states. Therefore, we have employed an integrated strategy encompassing nuclear magnetic resonance spectroscopy, mutagenesis, biochemistry, and mass spectrometry. Our studies reveal a unique thioester-thiol exchange mechanism in IacP, which transfers the acyl chain from the phosphopantetheine arm to a reactive cysteine in loop II, located distal to the 4’PP attachment site. To our knowledge, such an unusual intramolecular acyl chain relay reaction has not been reported previously for any other ACP. Notably, the reactive cysteine (Cys 54) lies within the SipB interaction interface of IacP, suggesting that this acyl-chain relay phenomenon may be directly linked to IacP-mediated host invasion and *Salmonella* virulence.

## Materials and methods

### Cloning, expression, and purification

The *Salmonella* Typhimurium IacP gene (UniProtKB P0CL00) was cloned into a pET28a (+) vector using Nde I and Hind III restriction sites to generate an N-terminal His-tagged fusion protein. The resulting plasmid was transformed into *E. coli* BL21 Rosetta (DE3) cells and protein expression was induced at 25 °C using 0.5 mM IPTG, followed by overnight incubation at 25 °C with constant shaking at 180 rpm. The cells were pelleted by centrifugation at 5,000 g at 4 °C. The supernatant was discarded, and the pellet was stored at −20 °C.

For purification, the frozen cell pellets were thawed, centrifuged at 20,000 g at 4 °C, and the supernatant was loaded onto a Ni^2+^-NTA affinity chromatography column, pre-equilibrated with 50 mM Tris-HCl, pH 7.8, and 150 mM NaCl. Eluted pure fractions were pooled, buffer exchanged to remove imidazole, and concentrated. The His tag was subsequently cleaved using immobilized thrombin. Final purification was achieved by passing the sample through an AKTA Superdex 75 size-exclusion chromatography column.

For *in vitro* assays, Sfp (a 4’-Phosphopantetheinyl transferase from *Bacillus subtilis*) was expressed separately. The protein was expressed using 0.5 mM IPTG at 25 °C, followed by overnight incubation, with constant shaking at 180 rpm. The cells were pelleted and stored at −20 °C. The next day, the pellet was thawed and sonicated. The supernatant was enriched using 30-60% ammonium sulfate precipitation, followed by desalting on a PD-10 column. The desalted lysate was subjected to anion-exchange chromatography (Akta Resource Q column) preequilibriated with 20 mM Tris-HCl, pH 8.0, 10 % glycerol, and eluted with a linear NaCl gradient from 0 to 1 M in the same buffer. Fractions were analyzed on a 12% SDS-PAGE, and the relatively pure fractions containing Sfp were pooled and loaded onto a Superdex 75 AKTA gel filtration column equilibrated with 20 mM Tris-HCl, pH 8.0, 150 mM NaCl, and 10 % glycerol. Eluted fractions were once again analyzed on a 12% SDS-PAGE, and the fractions exhibiting homogeneity were pooled, concentrated, and stored in 50% glycerol at −20 °C.

### Expression in minimal media

To produce uniformly labeled ^15^N^13^C IacP and its mutants, a single *E. coli* Rosetta (DE3) colony harboring the pET28a (+) plasmid encoding IacP was cultured in 5 ml of LB medium. Approximately 1% of the overnight culture was used to inoculate 1 liter of M9 (minimal) media supplemented with thiamine HCl, vitamins, and antibiotics. Calcium chloride and magnesium sulfate were added to a final concentration of 0.1 mM and 1 mM, respectively. Protein expression was induced by adding 0.5 mM IPTG when the culture’s absorbance at 600 nm reached 0.8-1.0, followed by overnight incubation (16 h) at 25 °C with constant shaking at 180 rpm. The bacterial cells were harvested by centrifugation at 5,000 g at 4 °C, and the pellets were stored at −20 °C. Subsequent purification steps were performed analogously to the unlabeled protein.

### Mutagenesis studies

IacP mutants were generated using a site-directed mutagenesis approach that utilizes Pfu Turbo (Agilent) and complementary oligo primers. The mutations were confirmed by Sanger DNA sequencing.

### *In vitro* Sfp assay to convert apo-to holo-/acyl-IacP^N^ intermediates

Acyl-IacP^N^ intermediates were prepared using apo-D35NIacP (referred to as apo-IacP^N^), in combination with acyl-CoAs of different lengths, and Sfp, a 4’-Phosphopantetheinyl transferase from *Bacillus subtilis*. A standard enzyme assay was carried out in a 2 ml reaction volume comprising 20 µM IacP^N^ (or mutants), 50 mM Tris-HCl (pH 8.0), 5 mM MgCl_2_, 5 µM Sfp, and 100 µM Coenzyme A or acyl-CoA of varying chain lengths. For NMR samples, the assay was scaled up proportionally.

Reactions were incubated at 37 °C for 2 h, and the products were analyzed qualitatively on a 12% urea-PAGE (1.5 M urea) and quantitatively by ESI-MS. Prior to NMR data acquisition, the reaction mixtures were thoroughly washed with NMR sample buffer using a 3 kDa Centricon to remove salt and excess CoA and concentrated to the desired volume.

### NMR sample preparation and data acquisition

For two- and three-dimensional NMR experiments, ^15^N/^13^C-labeled 300 µM - 1 mM protein in 50 mM Tris-HCl, 150 mM NaCl, pH 7.8, and 10 % D_2_O (NMR sample buffer) was used. For DTT treatment, the wild-type IacP sample was incubated with 10 mM DTT for 2h at 40°C in the NMR sample buffer before data acquisition. For alkylation/iodoacetamide (IAA) treatment, apo-IacP^N^ was incubated with 10 mM IAA at room temperature in the dark for 2 hours, followed by thorough washing with the NMR sample buffer to remove excess IAA.

All NMR experiments were acquired at 293 K on a 700 MHz Bruker Avance III/Neo NMR spectrometer equipped with a TXI/QCI probe, installed at the National Institute of Immunology, New Delhi, India. Standard triple resonance experiments were acquired for the sequential backbone assignments of *Salmonella* Typhimurium IacP, *viz.* HNCACB, CBCAcoNH, HNCA, and CCcoNH experiments. The acquired raw data were subsequently processed on a workstation running Red Hat Enterprise Linux 5.0 using NMRPipe/NMRDraw(*37*). ^15^N/^13^C spectra were indirectly referenced using DSS as a chemical shift standard(*38*).

### Average amide chemical shift changes

Changes in the amide chemical shifts of the mutants have been reported as an average of proton and nitrogen chemical shifts (ΔΔδ_HN_) derived from Equation 1(*39*)

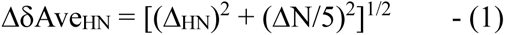

where Δ_HN_ is the change in the chemical shift in the proton dimension, and ΔN is the change in the nitrogen dimension.

### ESI-MS analysis

Due to the hydrophobic nature of IacP, the samples were desalted using C4 ZipTips. The ZipTips were first preconditioned with a solution containing 0.1% formic acid in 100% acetonitrile, then with a solution containing 0.1% formic acid in 50% acetonitrile (twice), and finally with 0.1% formic acid in autoclaved water (twice). After this, a 20 μl protein sample was repeatedly aspirated and dispensed through the ZipTip (>10 cycles). The unbound protein was removed with 0.1% Formic Acid in 5% Methanol. The protein bound to the ZipTip was eluted in 20 μl 50 % acetonitrile containing 0.1 % formic acid. The eluted sample was loaded onto a Thermo Scientific Orbitrap Exploris/Q Exactive Plus and processed using Xcalibur software. The intact protein masses were determined by deconvoluting the observed charge-state distribution.

## Results

### The amino acid sequence of *Salmonella* invasion acyl carrier protein (IacP) is distinct from that of its FAS AcpP

The *Salmonella* Typhimurium genome encodes two distinct acyl carrier proteins: the canonical AcpP (UniProtKB P0A6B1), which functions in the fatty acid synthesis (FAS) pathway, and the noncanonical invasion acyl carrier protein (IacP, UniProtKB P0CL00), which is indispensable for *Salmonella* entry into the intestinal epithelial cells. The two ACPs share 29% sequence identity, as illustrated in Fig. 1A. The conserved ^34^ADSL^37^ motif, crucial for the recognition of carrier proteins by the Holo-acyl-carrier protein synthase (AcpS), is conserved, as both serve as substrates for *Salmonella* AcpS. Besides, several other residues, crucial for interactions with FAS enzymes or AcpP function, are markedly different in IacP. For instance, Phe 28 present near the opening of the hydrophobic cavity of AcpP is substituted with a leucine, Thr 39 with a leucine, Met 44 with a Phe, Ala 45 with a Gly, Thr 52 with Ile, Ile 54 with a Cys, Pro 55 with an Asn, Ala 59 with a Leu, and Thr 63 with a methionine, and several others. Apart from Cys 54 present in loop II, two more free cysteines are present in IacP, at position 14 (end of helix I) and 69 (helix IV). Notably, most type II FAS ACPs lack a cysteine, most likely to avoid interaction with the phosphopantetheine arm. Interestingly, the atypical type II ACPs of *Salmonella* and non-ribosomal peptide synthetase carrier proteins (PCPs) harbor one or more cysteines, whose function remains unclear. The sequence comparison of IacP with the ACPs associated with the type III secretion system (T3SS) from other organisms displays low conservation, suggesting differences in their mechanism of function (Fig. 1B).

**Fig. 1.**
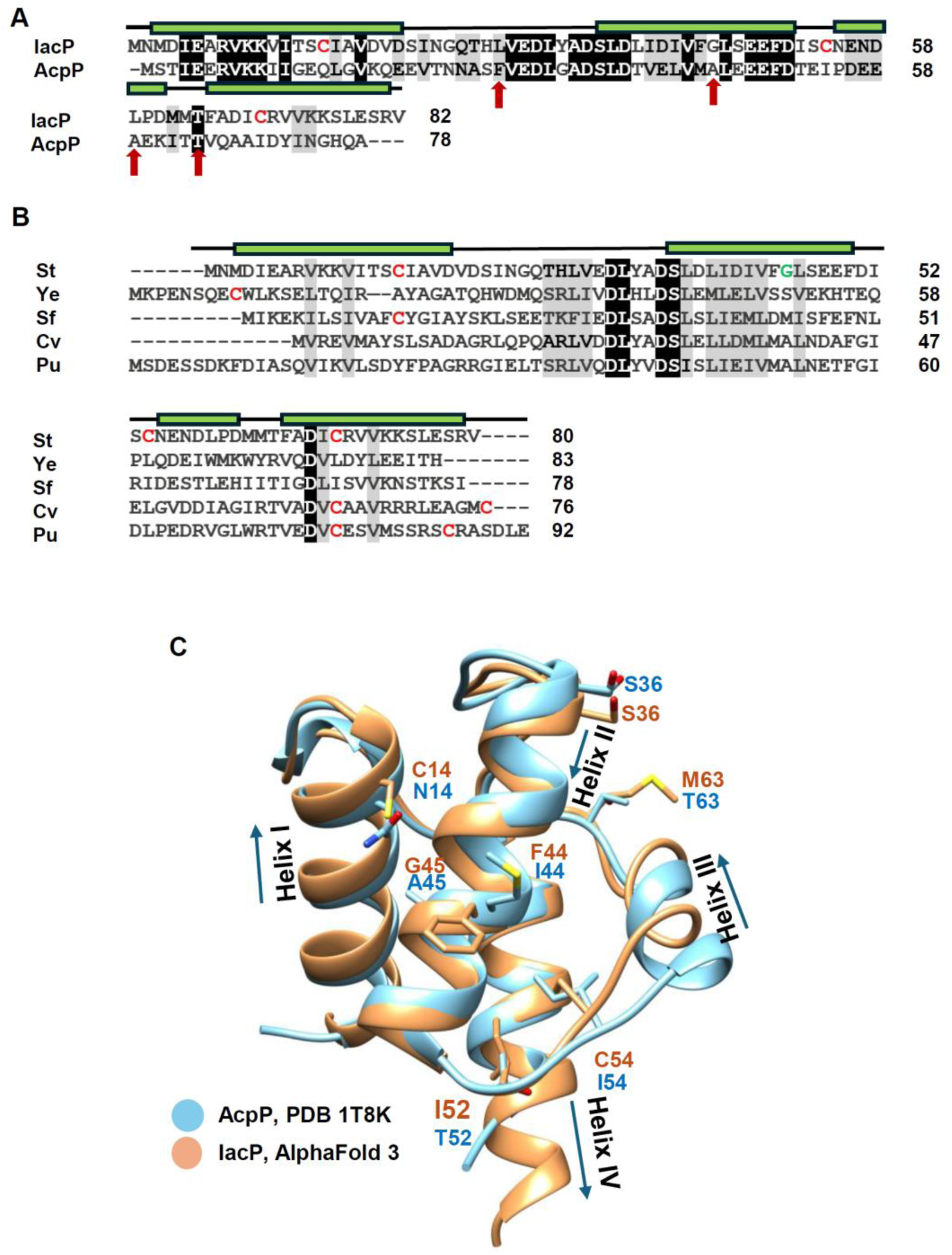
Sequence comparison of *Salmonella* Typhimurium IacP with its AcpP and other noncanonical ACPs involved in the type III secretion system. A) Alignment of the *Salmonella* IacP sequence with that of AcpP (UniProtKB P0A6A8). B) Multiple sequence alignment of IacP with representative noncanonical ACPs involved in the type III secretion system. C) Structural superposition of *E. coli* AcpP PDB 1T8K (sequence identical to *Salmonella* AcpP) onto the AlphaFold 3 predicted model of *Salmonella* IacP. Abbreviations used-St; *Salmonella* Typhimurium (UniProtKB P0CL00), Ye; *Yersinia enterocolitica* subsp. *enterocolitica* 8081 (NCBI WP_050139102.1), Sf; *Shigella flexneri* M90T (NCBI EFY8639960.1), Cv; *Chromobacterium violaceum* ATCC 12472 (UniProtKB Q7NUT2), Pu; *Pseudomonas* sp. UW4 (NCBI WP_015096039.1).

Notably, *E. coli* AcpP sequence is 100% identical to *Salmonella* Typhimurium FAS AcpP. Consequently, the *E coli* AcpP crystal structure (PDB 1T8K) has been used as a representative of *Salmonella* Typhimurium AcpP in this study. In the absence of an experimental IacP structure, the AlphaFold 3 predicted model has been employed for structural analysis (*40*). Both *Salmonella* AcpP and IacP display a four-helical bundle fold comprising helices I-IV, with helix III as a 3_10_ helix, as in most type II FAS ACPs. For consistency, we have used the *E. coli* AcpP (PDB 1T8K) residue numbering for IacP, placing Ser at position 36 in the ‘DSL’ motif. As illustrated in Fig. 1C, the crystal structure of *Salmonella* AcpP, shown as ribbons (PDB 1T8K, sky blue), is superimposed on the *Salmonella* IacP structure (AlphaFold 3 model, colored tan)(*40*). Residues important for AcpP function that differ between the two proteins are depicted as sticks and colored by heteroatom type, with the conserved Ser 36 side chain highlighted in both structures. Overall, the two proteins display a similar fold, though helix IV of IacP is slightly extended, with an extra turn at the C-terminus.

### The acylated forms of IacP display two sets of peaks in the ^1^H^15^N HSQC spectrum

To probe the molecular mechanism of acyl chain sequestration by IacP, acyl intermediates were generated *in vitro* using Sfp and a series of acyl-CoAs. Since the wild-type IacP is expressed exclusively in the holo-form in *E. coli,* a D35NIacP mutant (referred to as IacP^N^ hereafter) was generated to express the protein in the apo-form (*41*). In a previous study from our group, we have shown that IacP^N^ expresses as an apo-protein in *E. coli* (*41*). The backbone chemical shifts of apo-IacP^N^ were assigned based on previously published assignments of wild-type holo-IacP and further validated using standard three-dimensional NMR experiments, including HNCACB, CBCAcoNH, HNCA, and CCCcoNH. The majority of the assignments were in good agreement with the previously reported values (*42*). Amide resonances corresponding to residues ^54^CNE^56^ could not be obtained, likely due to conformational/chemical exchange. The ^1^H^15^N HSQC spectrum of apo-IacP^N^ (apo-D35NIacP, colored green) superimposed on the S36AIacP (colored red) is shown in Fig. S1, with all the peaks labeled (Fig. S1). Minor changes were observed near the sites of mutation (D35 and S36).

To elucidate the mechanism of chain sequestration by IacP, *in vitro* apo-IacP^N^ was converted to holo-IacP^N^ and various acyl-IacP^N^ intermediates using Sfp (a 4’-phosphopantetheinyl transferase with broad substrate specificity), and the conversions were verified by 12% urea-PAGE (Fig. S2A). A clear change in the electrophoretic mobility was observed upon formation of holo- and acyl-IacP^N^ intermediates, attributable to the gain of a negative charge upon attachment of the phosphopantetheine arm (PO_4_^-^). All the acyl-IacP^N^ intermediates displayed a very similar migration on urea-PAGE, akin to that of holo-IacP^N^ (Fig. S2A). The successful formation of the acyl intermediates was further confirmed by ESI-MS analysis (Fig. S2 A-J). In the figure, the first panel for each intermediate displays the raw data, and the second panel shows the deconvoluted envelope. The expected and determined masses were within ±1 Da of each other in each case.

Figure 2 displays selected regions of a multiple overlayed ^1^H^15^N HSQC spectra for C4- (magenta), C6- (pink), C8- (green), C10- (orange), C12- (blue), C14- (cyan), and C16-IacP^N^ (black) intermediates, superimposed on the holo-IacP^N^ (red) spectrum, highlighting four amides in particular, *viz.* Leu 28, Gly 45, Leu 59, and Thr 64, that exhibit pronounced chemical shift perturbations upon acylation. Notably, three of these residues (Leu 28, Gly 45 & Leu 59) differ from their amino acid counterparts in AcpP (shown by an upward-pointing red arrow in Fig. 1A).

**Fig. 2.**
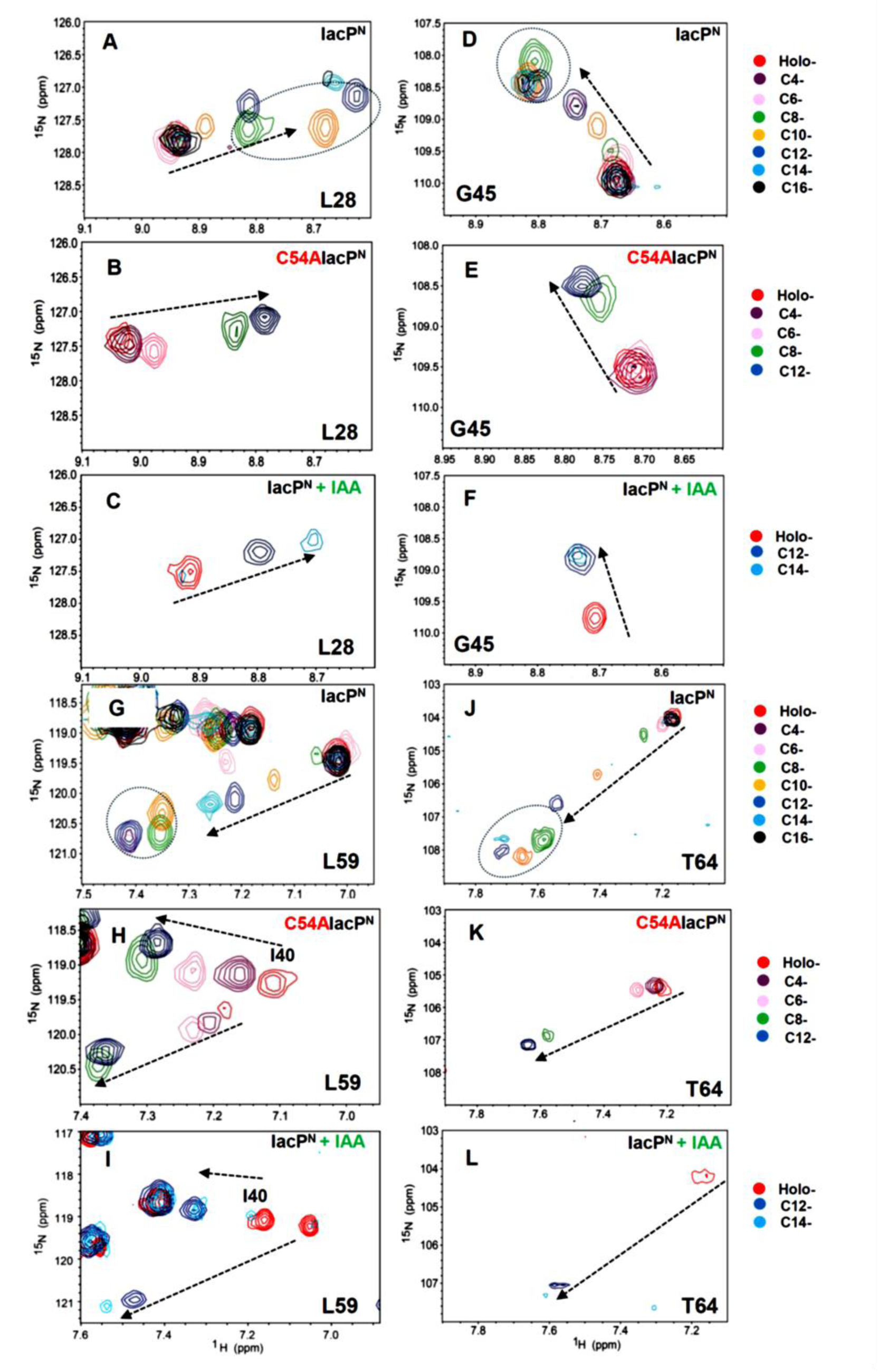
The acyl-IacP^N^ intermediates display two distinct sets of resonances in the ^1^H^15^N HSQC spectra, while the C54AIacPN intermediates display only one set of peaks. Selected regions of the ^1^H^15^N HSQC spectra displaying peaks for holo- (red), C4- (magenta), C6- (pink) C8- (green), C10- (orange), C12- (dark blue), C14- (cyan) and C16- (black) IacP^N^ intermediates, for **A)** Leu 28, **D)** Gly 45, **G)** Leu 59, and **J)** Thr 64 amides. The peaks shown in circles highlight the Cys-bound peaks, while those that show a change in chemical shift with increasing acyl-chain length are 4’PP-bound, indicated by an arrow. Selected regions of a superimposed ^1^H^15^N HSQC spectra displaying peaks for holo- (red), C4- (magenta), C6- (pink), C8- (green), and C12- (blue) C54AIacP^N^ intermediates for **B)** Leu 28, **E)** Gly 45, **H)** Leu 59, and **K)** Thr 64 amides. Only one set of peaks corresponding to the 4’PP-bound IacP^N^ peaks was visible in the intermediates of the C54A mutant. Dashed arrows indicate the direction in which chemical shifts change. Superposition of holo- (red), C12- (blue), and C14- (cyan) S-carboxyamidomethyl-IacP^N^ spectra displaying peaks for **C)**, Leu 28, **F)** Gly 45, **I)** Leu 59, and **L)** Thr 64 amides. Only the 4’PP-bound peaks were visible in this covalently modified form of IacP^N^ as well. The color coding used for each intermediate is labeled on the right side of the figure.

Unexpectedly, in the ^1^H^15^N HSQC spectra of C6-, C8-, C10-, and C12-IacP^N^ intermediates, two distinct resonances were observed for each of the amides (Fig. 2A, D, G & J). The peak positioned proximal to the holo-peak displayed a consistent change in chemical shift with increasing acyl-chain length as we go from holo-(red) to C12-(cyan), akin to the 4’PP-bound conformation of canonical type II ACPs (*43*). Therefore, the peak was assigned as the 4’PP-bound state. This 4’PP bound peak was less intense for the C6- and C8-conformers, but for longer chains it was relatively more intense. The direction of chemical shift change of this peak with increasing acyl-chain length is indicated by a black arrow. In contrast, a second peak, previously uncharacterized, is located farthest from the holo-peak, but closer to the C12-4’PP-bound IacP^N^ conformation. This second peak displayed a significantly larger peak volume and was invariant across all acyl intermediates (C6-C12-IacP^N^), suggesting insensitivity to changes in acyl chain length. The uncharacterized peak is shown by a circle in the figure.

The two conformers (4’PP & uncharacterized) were observed for several of the amides of the acyl-IacP^N^ intermediates, Leu 28, Val 29, Glu 30, Leu 32, Asn 35, Leu 37, Ile 40, Ile 42, Gly 45, Leu 46, Ile 52, Leu 59, Thr 64, and Cys 69. In Fig. S4A, the amides that exhibit peak twinning in the acyl intermediates of IacP^N^ in the ^1^H^15^N HSQC spectrum have been mapped onto the ribbon representation of the AlphaFold 3 model of IacP and highlighted in green.

We next compared the Cα chemical shifts of the C8-(green) and C10-IacP^N^ (brown) intermediates by analyzing their HNCA spectra, using the holo-IacP^N^ (red) spectra as a reference. Some residues, namely Val 7, His 27, Ile 40, Thr 64, and Ile 68, displayed noticeable Cα chemical shift change. These residues have been mapped onto the ribbon representation of IacP in Fig S4B, and highlighted in green. Regions of the ^13^C^15^N HNCA spectra displaying the corresponding peaks are shown in Fig. S5A-F. Of the five peaks displaying noticeable Cα chemical shift change, the largest change, ∼2 ppm, was observed in the Thr 64 chemical shift (Fig. S5E).

### The acyl-C54AIacP^N^ intermediates demonstrate that the second conformation arises due to the interaction of Cys 54 with the acyl chain

To probe the role of cysteines in giving rise to the second set of conformations in the acyl-IacP^N^ intermediates, cysteine mutants, i.e., C14A, C54A, and C69A IacP^N^, were generated and enzymatically converted into the acyl-IacP^N^ intermediates. Interestingly, the C54AIacP^N^ intermediates (C4-, C6-, C8-, C10-, and C12-) displayed only one set of peaks in the ^1^H^15^N HSQC spectrum, shown for some of the representative amides, *i.e.,* Leu 28, Gly 45, Leu 59, and Thr 64, in Fig. 2B, E, H, and K, respectively. Acylation was confirmed by urea-PAGE (shown in Fig. S2B) and ESI-MS analysis (shown in Fig. S3 K-P). All the peaks of the acyl-C54AIacP^N^ intermediates displayed a systematic chemical shift change with increasing acyl chain length, and their chemical shifts aligned well with the 4’PP-bound conformation of acyl-IacP^N^ intermediates in the ^1^H^15^N HSQC spectra. The absence of the second set of peaks in the acyl-C54AIacP^N^ intermediates strongly supported the role of Cys 54 side chain interaction with the acyl chain in giving rise to the uncharacterized peak.

### The acyl-S-carboxyamidomethyl-IacP^N^ intermediates also display a single conformational state in the ^1^H^15^N HSQC spectrum

As an alternative strategy to validate the role of Cys 54 in generating the Cys-bound conformation in the acylated-IacP^N^ intermediates, the cysteine side chains of IacP^N^ were covalently modified with iodoacetamide. The S-carboxyamidomethyl-IacP^N^ was prepared by incubating apo-IacP^N^ with 10 mM iodoacetamide (IAA). This sulfhydryl alkylating agent irreversibly alkylates cysteine thiols. Superposition of the S-carboxyamidomethyl-IacP^N^ on the apo-IacP^N^ ^1^H^15^N HSQC spectra revealed minor chemical shift perturbations, suggesting that alkylation does not significantly alter the overall protein fold (data not shown).

Next, the S-carboxyamidomethyl-IacP^N^ was converted into holo-, C12-, and C14-intermediates. The C12 modification was verified by ESI-MS (Fig. S3 O & P). In the ESI-MS spectra shown, equimolar amounts of C12-S-carboxyamidomethyl-IacP^N^ and C12-C54AIacP^N^ were mixed together and loaded. The determined masses were consistent with a C12-S-carboxyamidomethyl-IacP^N^ intermediate in which only one cysteine has been modified and a C12-C54AIacP^N^ intermediate. Conceivably, the other two cysteines are not fully exposed and therefore they do not get modified. A small amount of holo-peak corresponding to the S-carboxyamidomethyl-IacP^N^ and C54AIacP^N^ was also observed due to the hydrolysis of the C12-intermediate. The ESI-MS spectrum of the C14-S-carboxyamidomethyl-IacP^N^ sample could not be obtained due to its rapid rate of hydrolysis and a tendency to aggregate.

Notably, the ^1^H^15^N HSQC spectra of C12- (blue) and C_14_- (cyan) S-carboxyamidomethyl-IacP^N^ intermediates also displayed a single set of peaks for Leu 28, Gly 45, Leu 59, and Thr 64 amides, as shown in Fig. 2C, F, I, and L, respectively. The position of these peaks in the ^1^H^15^N HSQC spectrum suggests that it is present in the 4’PP-bound conformation, further supporting the conclusion that the second conformation arises due to the transfer of the acyl-chain to a distal cysteine (Cys 54).

### The acyl- chain is sequestered in the core of the 4’PP-bound acyl-C54AIacP^N^ conformers

Since the ^1^H^15^N HSQC spectra of the acyl-C54AIacP^N^ intermediates displayed a single set of peaks corresponding to the 4’PP-bound conformation of IacP^N^, their amide chemical shift changes were monitored to understand the mechanism of chain sequestration in the 4’PP-bound conformation. A very similar overall pattern of perturbation was observed for C6-, C8-, and C12-C54AIacP^N^ intermediates. However, with increasing acyl chain length, the magnitude of perturbations increased as expected in the 4’PP-bound intermediates, shown in Fig. 3B-D. In the figure, a discontinuous horizontal threshold indicates one standard deviation. Of all the amides, Thr 64 displayed the largest perturbations, with AveHN > 0.5 ppm. Nevertheless, significant chemical shift perturbations (>1 SD) were observed for residues present in all four helices. The average amide chemical shift changes upon formation of the C12-C54AIacPN intermediate have been mapped onto the AlphaFold 3 model of IacP, shown in Fig 3E, shown as worms. The swellings on the worm are directly proportional to the magnitude of the amide chemical shift perturbations observed. In the figure, all four helices and loops exhibit chemical shift changes as acyl chain length increases, confirming the chain’s sequestration in the hydrophobic core. The C4-intermediate also displayed minor perturbations, with a pattern similar to that of the other intermediates, suggesting partial sequestration (Fig. 3A).

**Fig. 3.**
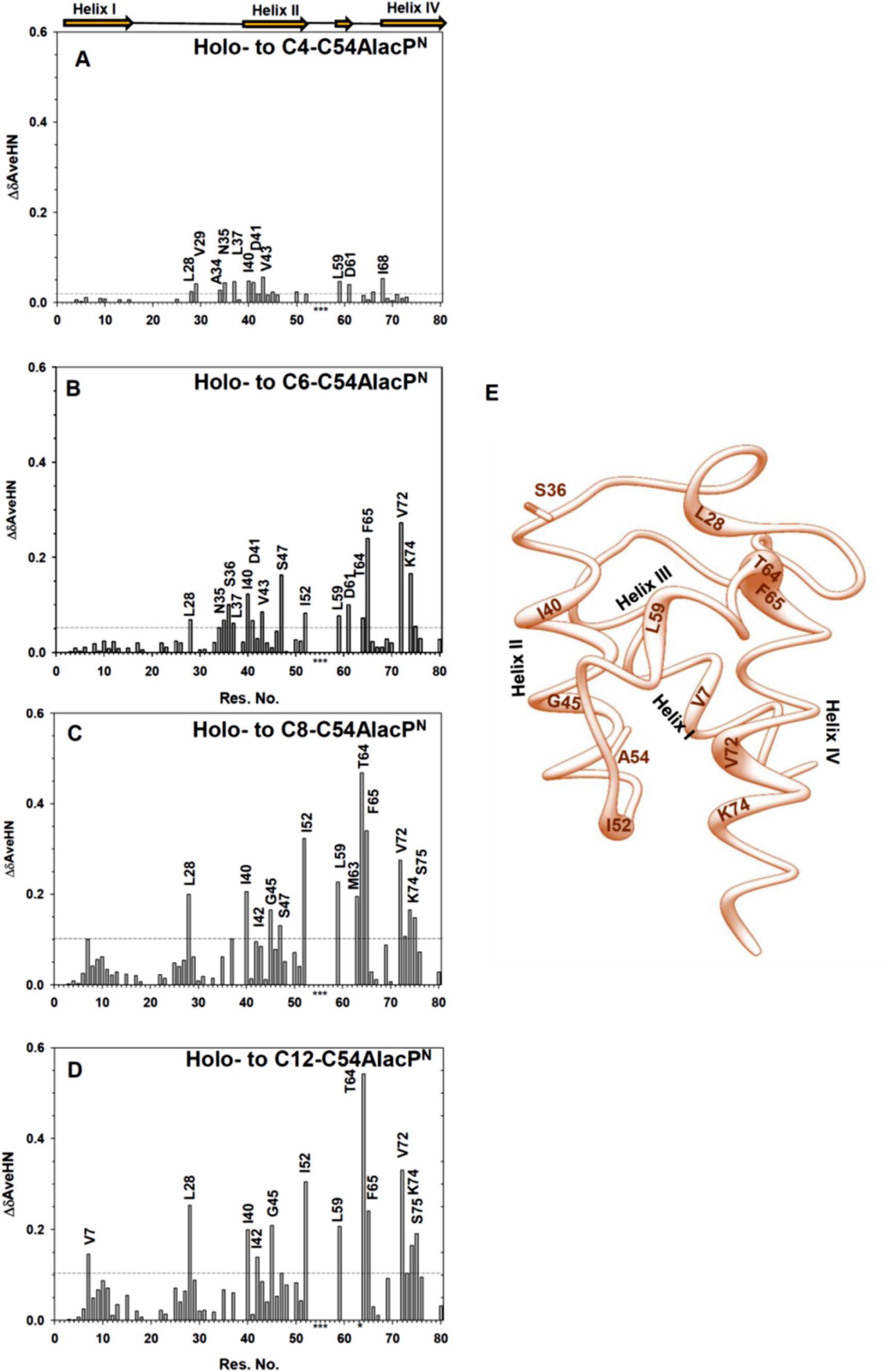
The amides displaying perturbations in the C54AIacP^N^ intermediates reside in loop I-helix IV. Average amide chemical shift changes are shown for the conversion of holo-C54A IacP^N^ to **A)** C4-, **B)** C6-, **C)** C8-, and **D)** C12-C54A IacP^N^ intermediates as a function of residue number. **E)** Average amide chemical shift changes observed upon conversion of holo- to C12-C54A IacP^N^ have been mapped onto the AlphaFold 3 model of IacP. The thickness of the worm at each point is directly proportional to the magnitude of the chemical shift change.

### The Cys-bound IacP^N^ conformer is more stable than the 4’-PP-bound IacP^N^ conformer and has a more expanded core

Correlation of the ^1^H and ^15^N chemical shifts of the cysteine-bound (acyl-Cys) and 4’PP-bound conformers in the various acyl-IacP^N^ intermediates is shown in Fig. 4A-D, where the Cys-bound peaks (the new peak observed in the IacP^N^ intermediates) are colored red, while the 4’PP-bound peaks are colored black. The 4’PP-bound peaks (black) displayed a systematic progression with increasing acyl chain length. In contrast, the Cys-bound peaks (red) were clustered within a narrow region, closely aligning with the chemical shifts of the C12- or C14-4’PP-bound IacP^N^ intermediates.

**Fig. 4.**
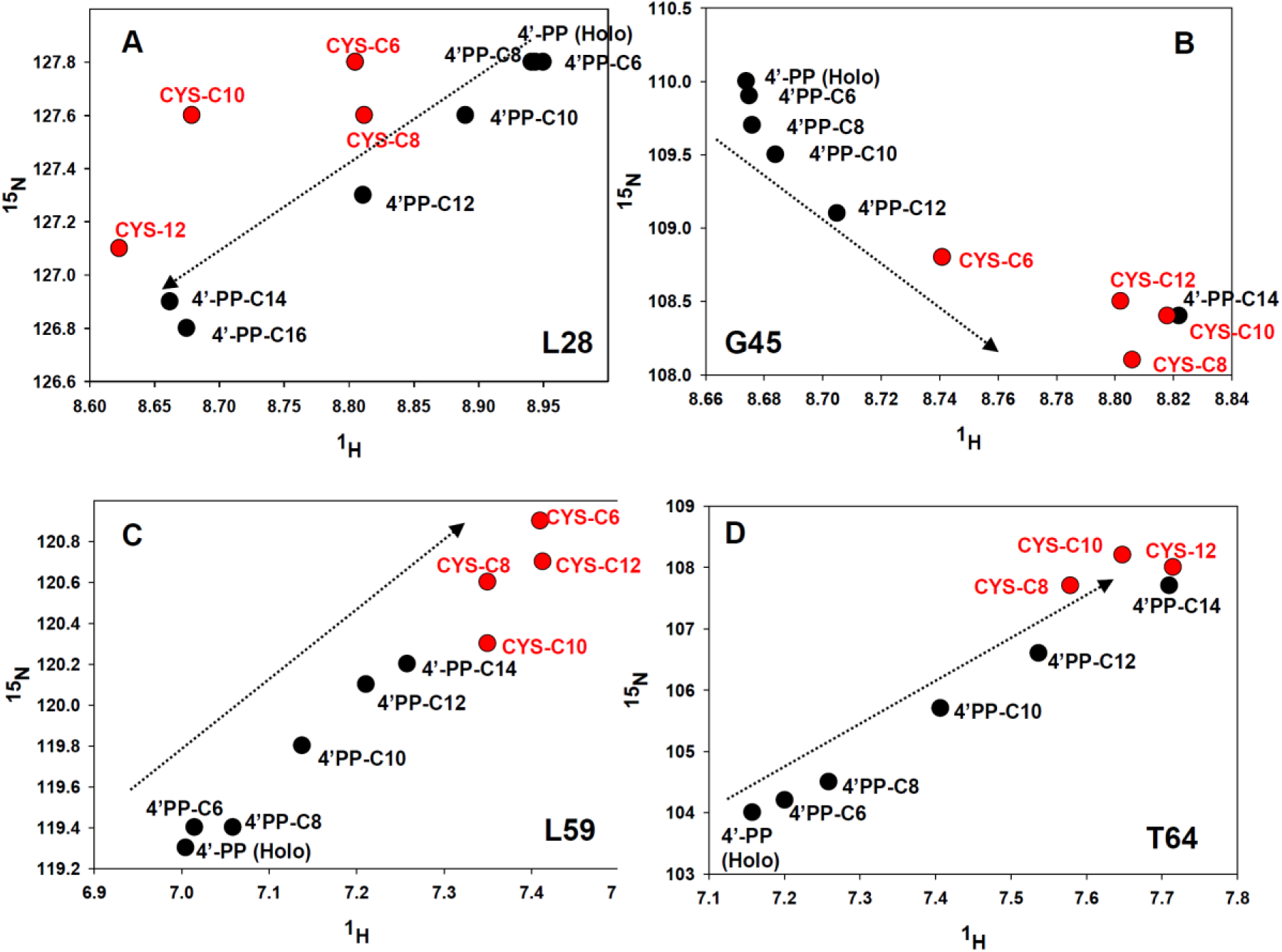
The Cys-bound IacP^N^ conformer has a more expanded core. ^1^H and ^15^N backbone correlations for the 4’PP- (black) and Cys-bound (red) conformers observed in the IacP^N^ intermediates C4-, C6-, C8-, C10-, C12-, C14-, and C16- IacP^N^ intermediates for residues **A)** Leu 28, **B)** Gly 45, **C)** Leu 59, and **D)** Thr 64. The Cys-bound conformers of all intermediates are clustered together, whereas the 4’PP-bound conformers exhibit a progressive change with increasing acyl chain length.

The trend was consistent in all four amides, Leu 28, Gly 45, Leu 59, and Thr 64. For instance, the C6-Cys and C8-Cys bound peaks (colored red) of Leu 28 resonate at ∼8.83 ppm, very close to the C12-4’PP-bound conformation. Similarly, the C10- and C12-Cys-bound conformations align with the C14-/C16-4’PP-bound peaks for Leu 28 (Fig. 4A). Likewise, in Gly 45, Leu 59, and Thr 64 amides (Fig. 4B-D), the Cys-bound conformer for C6-, C8-, C10-, and C12-IacP^N^ were clustered at a very similar ^1^H chemical shift value, close to the ^1^H chemical shift value of the C14-4’PP-bound conformer, suggesting a more expanded IacP^N^ core in the Cys-bound conformer. No peak twinning was observed in the C4-intermediate, and its chemical shift fully overlapped with the holo-peak.

The relative stability of the two conformational states was next assessed through a hydrolysis experiment. The breakdown of the 4’PP-bound (labeled P) and Cys-bound (labeled C) peaks of the C12-IacP^N^ was followed for 96 h at room temperature (20 °C), as shown in Fig. 5A-O. In the beginning of the experiment, both the peaks appear equally intense. Hydrolysis of Leu 28, Gly 45, and Leu 59 amides was followed in a ^1^H^15^N HSQC spectrum. The spectra revealed that the cysteine-bound conformers persisted with higher intensity and longer duration than the 4’PP-bound conformer, indicating greater stability of the Cys-bound conformer.

**Fig. 5.**
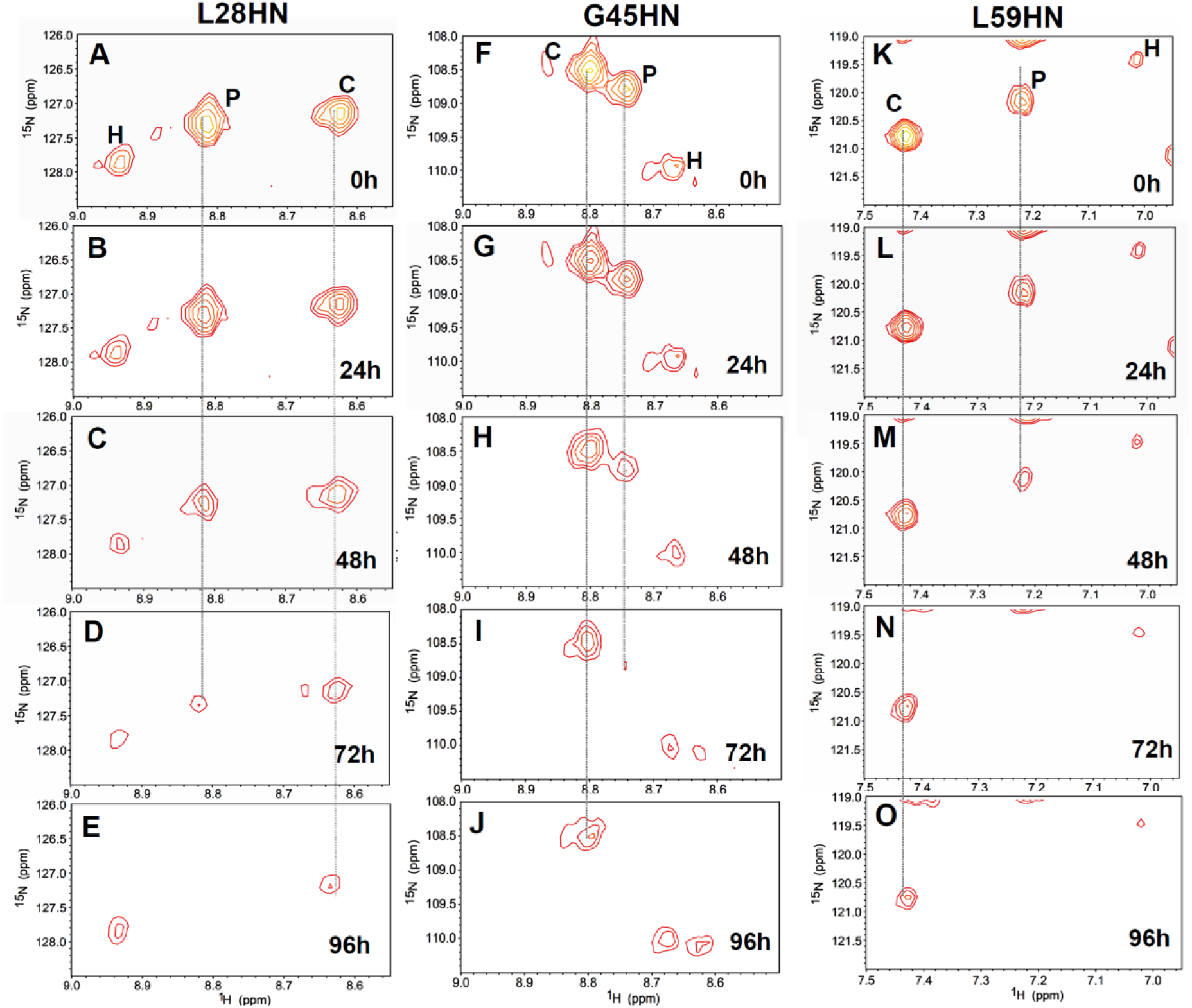
The Cys-bound IacP^N^ conformer is more stable than the 4’PP-bound conformer. Selected regions of the ^1^H^15^N HSQC spectra displaying changes in the peak volume as a function of time due to hydrolysis of the acyl-chain for the 4’PP-bound and Cys-bound conformers observed for Leu 28, Gly 45, and Leu 59 amides when the sample was left at 20 °C. No significant increase in holo-peak intensity was observed over time, as the protein tended to aggregate upon hydrolysis. The peak labeled C represents Cys-bound, P represents 4’PP-bound, and H represents holo-peak formed as result of acyl chain hydrolysis. The figure was prepared using Sparky(*44*).

## Discussion

The canonical acyl carrier proteins of the type II FAS pathway have been extensively characterized in *E. coli, Bacillus subtilis, Mycobacterium tuberculosis, Helicobacter pylori, Plasmodium falciparum, Leishmania major, Spinacea oleracea,* and several other organisms (*43, 45–55*). In contrast, the noncanonical ACPs still remain underexplored. To fill this gap in our understanding of atypical ACPs, a comprehensive study was undertaken of their acyl intermediates. Our results demonstrate that the noncanonical *Salmonella* IacP diverges from its type II FAS AcpP in several respects. It differs in sequence from the type II FAS ACP across multiple sites in the protein. An important and evolutionarily conserved residue of most FAS AcpPs is Ile 54, which is critical for the structure and function. This residue is substituted with a cysteine in IacP (*56*). An Ile is present at position 52 instead, which may partially compensate for the loss of Ile 54. Gly 45 is another interesting residue in helix II of IacP, which is an alanine in AcpP. A glycine in a helix can introduce kinks, which might be important for IacP dynamics and function(*57*). Structurally, both AcpP and IacP have a conserved four-helix bundle fold. The first three helices of IacP (helices I-III) and AcpP are comparable; however, helix IV is extended by an extra turn in IacP, reminiscent of mitochondrial ACPs(*53*). Moreover, IacP appears to operate differently compared to the FAS AcpP. The ^1^H^15^N HSQC spectrum of the acyl-IacP^N^ intermediates displayed unique peak heterogeneity, markedly different from that of AcpP. Of the two peaks observed per amide, one corresponds to the 4’-phosphopantetheine (4’PP) bound conformation analogous to AcpP, and a second represents a previously uncharacterized peak. Interestingly, the acyl-C54AIacP^N^ & acyl-S-carboxyamidomethyl-IacP^N^ intermediates lacked the second peak in the ^15^N HSQC spectrum, demonstrating that the additional conformational state in the IacP^N^ intermediates arises from the transfer of the acyl-chain from the 4’PP arm to the free thiol of Cys 54.

The two conformational states of IacP^N^ intermediates- *i.e.,* the 4’PP- & the Cys-bound conformation exhibit pronounced structural, functional, and dynamic differences. The Cys-bound conformer displayed substantially larger amide chemical shift perturbations than the 4’PP-bound conformation, suggesting a more expanded IacP^N^ core. In addition, the Cys-bound conformer displayed a greater peak volume than the 4’PP-bound conformer for the acyl-chains upto C8-, indicating that the bulk of the IacP^N^ molecules efficiently relay the acyl chain from the 4’PP arm to the cysteine 54 side chain. Hydrolysis experiments further indicate that the Cys-bound acyl-IacP^N^ conformer is slightly more stable than the 4’-PP-bound conformer. This difference could arise due to the dynamic nature of the 4’PP-arm compared to the short side chain of cysteine that is directly attached to the acyl chain. Taken together, these findings highlight the high reactivity of Cys 54, which differentiates IacP from AcpP and explains IacP’s inability to complement FAS AcpP function in a temperature-sensitive mutant (*58*). A somewhat related thiol-exchange phenomenon was reported in a clinical context involving a heterozygous human variant of the type I FAS ACP (Arg2117Cys), in which the mutation of an arginine in loop II (at a similar position to Ile 54) to cysteine introduced a similar aberrant reactivity. In that patient, the mutant human fatty acid synthase (FASN) complex underwent non-enzymatic (chemical) acetyl group transfer to lysine *via* cysteine, triggering ubiquitination, subsequent proteosomal degradation, and ultimately FAS-deficiency-related pathologies (*59*). Together, these two independent studies highlight the role of context-dependent cysteine reactivity in mediating thioester exchange, which profoundly influences protein structure, stability, and function (*60*).

Since the HSQC spectra of the C54AIacP^N^ intermediates displayed only the 4’PP-bound peaks (conformation) with no acyl-exchange reaction with Cys 54, this mutant offered us an excellent opportunity to track the chemical shift changes of the 4’PP-bound conformer upon acyl chain sequestration. Akin to the FAS type II AcpP-where all four helices undergo significant perturbations during acyl chain elongation, the 4’PP-bound C12-C54AIacP^N^ intermediates also displayed perturbations at the amides of all four helices and loops, suggesting that the acyl chain is sequestered in the IacP^N^ core. However, unlike the type II FAS ACPs, in which the acyl chain reaches the bottom of the cavity and its tip interacts with Ile 54, in IacP, the interacting residue appears to be Ile 68 in IacP, based on the large perturbations observed for the Thr 64 amide & Cα, which is present at i-4 from Ile 68. Notably, Ile 64 is present at the beginning of helix IV and its backbone O is hydrogen bonded to Ile 68 HN. We compared the AlphaFold 3 IacP model with the AcpP structure of *E. coli* (PDB 1T8K), both of which contain a Thr at position 64. However, unlike IacP, this residue does not exhibit significant perturbations upon acylation in AcpP, further highlighting the differences in the mechanism of chain sequestration by the two proteins. In our studies, the Cys-bound conformer was observed only in the acyl-IacPN intermediates, where it appeared as a second peak alongside the 4’PP-bound conformer. The Cys-bound conformer displayed a similar pattern of perturbations as the 4’PP-bound conformer, and in the same direction, but displayed much larger changes compared to holo, consistent with a more expanded hydrophobic core.

Taken together, the present study provides novel insights into the molecular mechanism of IacP function and helps fill some of the gaps in our understanding of IacP-mediated SipB (*Salmonella* invasion protein B) maturation and, in turn, *Salmonella* infection. The pathogen relies on a Type III secretion system (T3SS) to infect the intestinal epithelial cells. SipB, a major translocon protein of the T3SS, serves as the acylation target (substrate) of IacP, a post-translational modification critical for the membrane localization of SipB, which marks the first step in the invasion of the host cells during *Salmonella* Typhimurium infection (*61, 62*). Palmitoylation of IacP on its phosphopantetheine arm has been shown to be critical for the catalytic function of IacP, and therefore, the S36A IacP mutant displays a loss-of-function phenotype (*5, 58, 61*). How an ACP like IacP could act as a catalyst and activate SipB is an open question. In that backdrop, the present work demonstrates a distinct acyl chain transfer mechanism in IacP, which shuttles the acyl chain moiety from the 4’PP arm to a distal cysteine (Cys 54). Notably, in our studies, this acyl-chain transfer was consistently observed for chain lengths from C6- to C12-IacP^N^. Since Cys 54 resides within the SipB interaction interface of IacP, as supported by NMR, we propose that the acyl chain transferred to Cys 54 of IacP may subsequently be relayed intermolecularly to Cys 231 of SipB (*42, 61*). Further *in vivo* studies are necessary to delineate the mechanistic details of this unique intermolecular acyl chain transfer during *Salmonella* infection. Most importantly, given that cysteine 54 is a critical residue mediating this acyl chain transfer, our studies point towards a potential therapeutic strategy: targeting cysteine 54 with cysteine-reactive inhibitors to impede *Salmonella* entry into the host cells.

### Accession IDs

- UniProtKB P0A6A8
- UniProtKB P0CL00
- NCBI WP_050139102.1
- NCBI EFY8639960.1
- UniProtKB Q7NUT2
- NCBI WP_015096039.1

## Data Availability Statements

The data supporting the findings of this study are available within the paper and in the Supporting Information. Additional files are available from the corresponding author on reasonable request.

## ACKNOWLEDGEMENTS

The genomic DNA of *Salmonella enterica* subsp. *enterica* 2004 Pennsylvania Tomato Outbreak, Serovar Typhimurium, was obtained through BEI Resources, NIAID, NIH: Isolate 1, NR 4614. The authors thank the Department of Biotechnology, Government of India, for financial and infrastructure support. The authors thank Dr. G. Senthil Kumar for his valuable suggestions during this study. Ms. Shanta Sen, Mass Spec Facility NII, is gratefully acknowledged for help with the acquisition of some of the ESI-MS data.

## ABBREVIATIONS

ACP: Acyl carrier protein
AcpP: *E. coli* acyl carrier protein
IacP: Invasion acyl carrier protein from *Salmonella* Typhimurium, PPTase, 4’Phosphopantetheinyl transferase
FAS: Fatty acid synthase
Sfp: 4’Phosphopantetheinyl transferase from *Bacillus subtilis*
AcpH: Holo-acyl carrier protein phosphodiesterase
SipB: *Salmonella* invasion protein B

