## Supplementary for "A thioester-thiol exchange reaction in *Salmonella* Invasion acyl carrier protein transfers the 4’PP-bound acyl chain to a distal cysteine": Fig S1.pdf

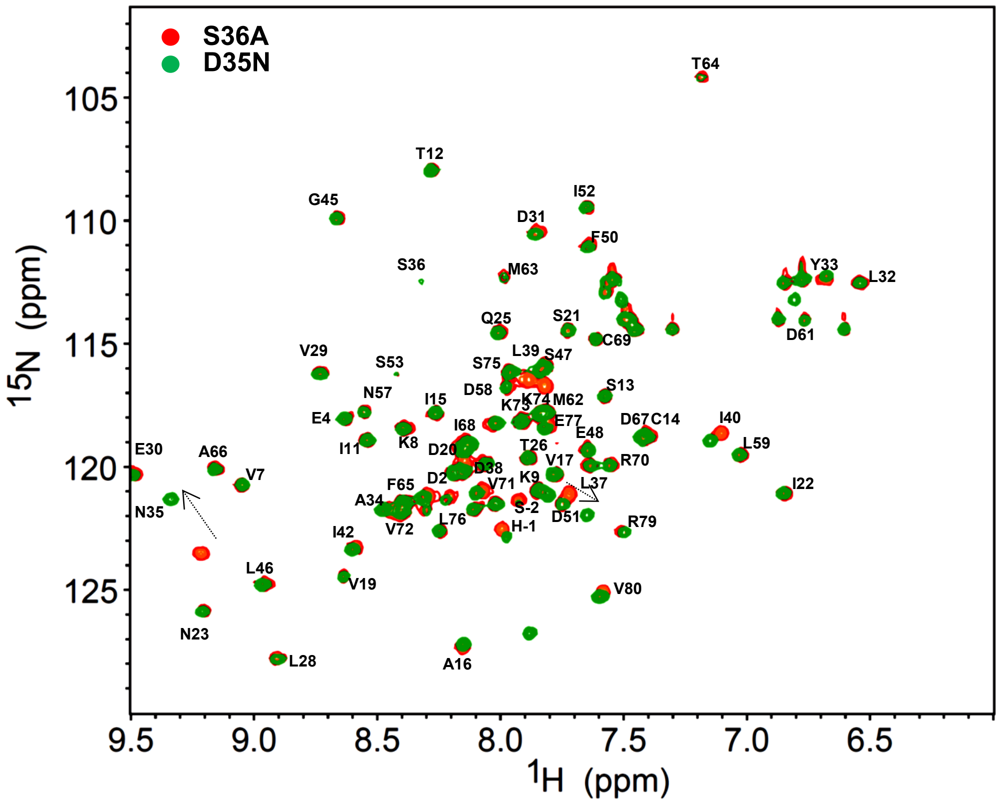


**Fig. S1 The D35N and S36A lacP mutants display a similar backbone conformation.** The $^{1}H^{15}N$ HSQC spectra of lacP$^{N}$ (D35N mutant, colored green) superimposed on the S36A lacP (colored red) demonstrate that they have a structurally similar backbone, except for changes observed near the site of mutation. The figure was prepared using Sparky [44].

A

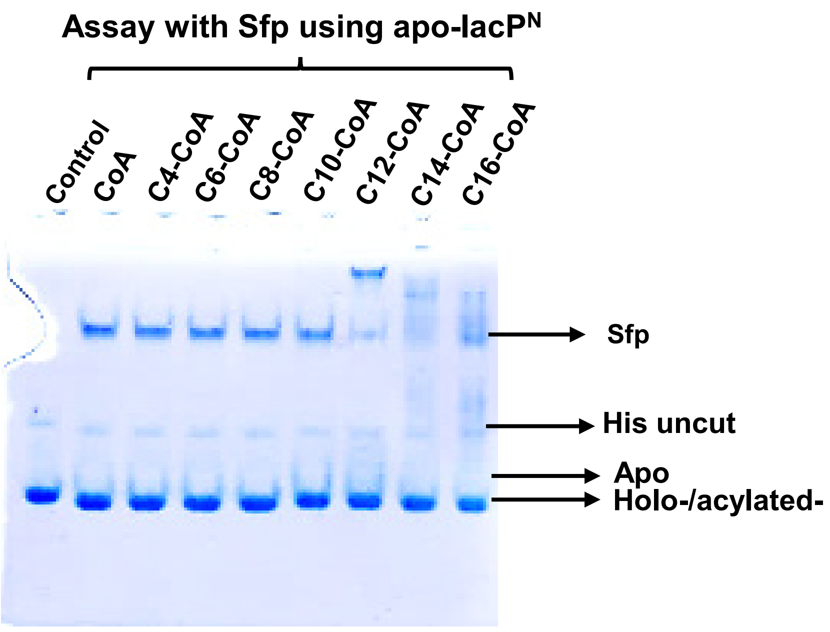


B

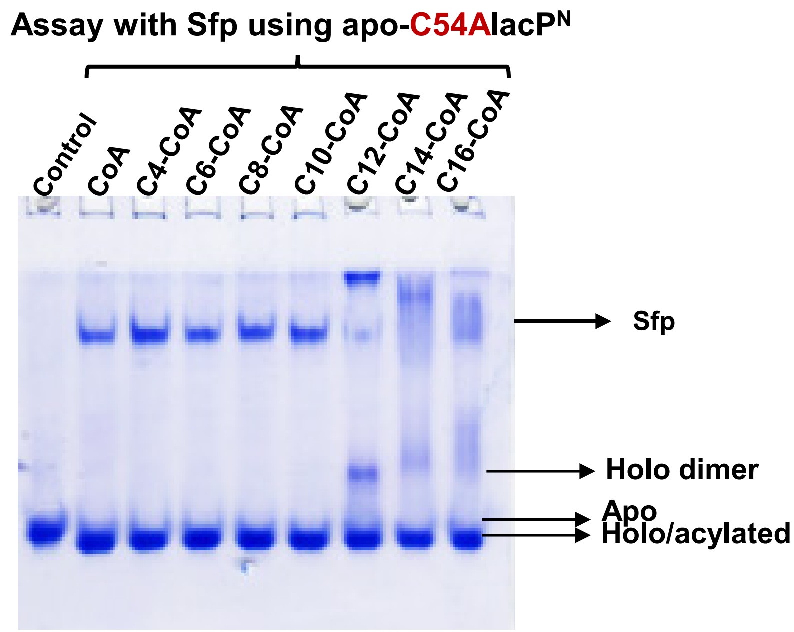


**Fig. S2 A 12% urea-PAGE (1.0 M urea) gel illustrating the conversion of apo- protein into a series of acylated intermediates.** Sfp assay for the conversion of A) apo-lacP$^{N}$, and B) apo-C54AlacP$^{N}$ into holo-, C4-, C6-, C8-, C10, C12-, C14-, and C16-acyl intermediates are shown. The acyl-CoAs used to generate the intermediates are labeled at the top of each lane. The bands corresponding to the apo- and holo-intermediates are labeled.

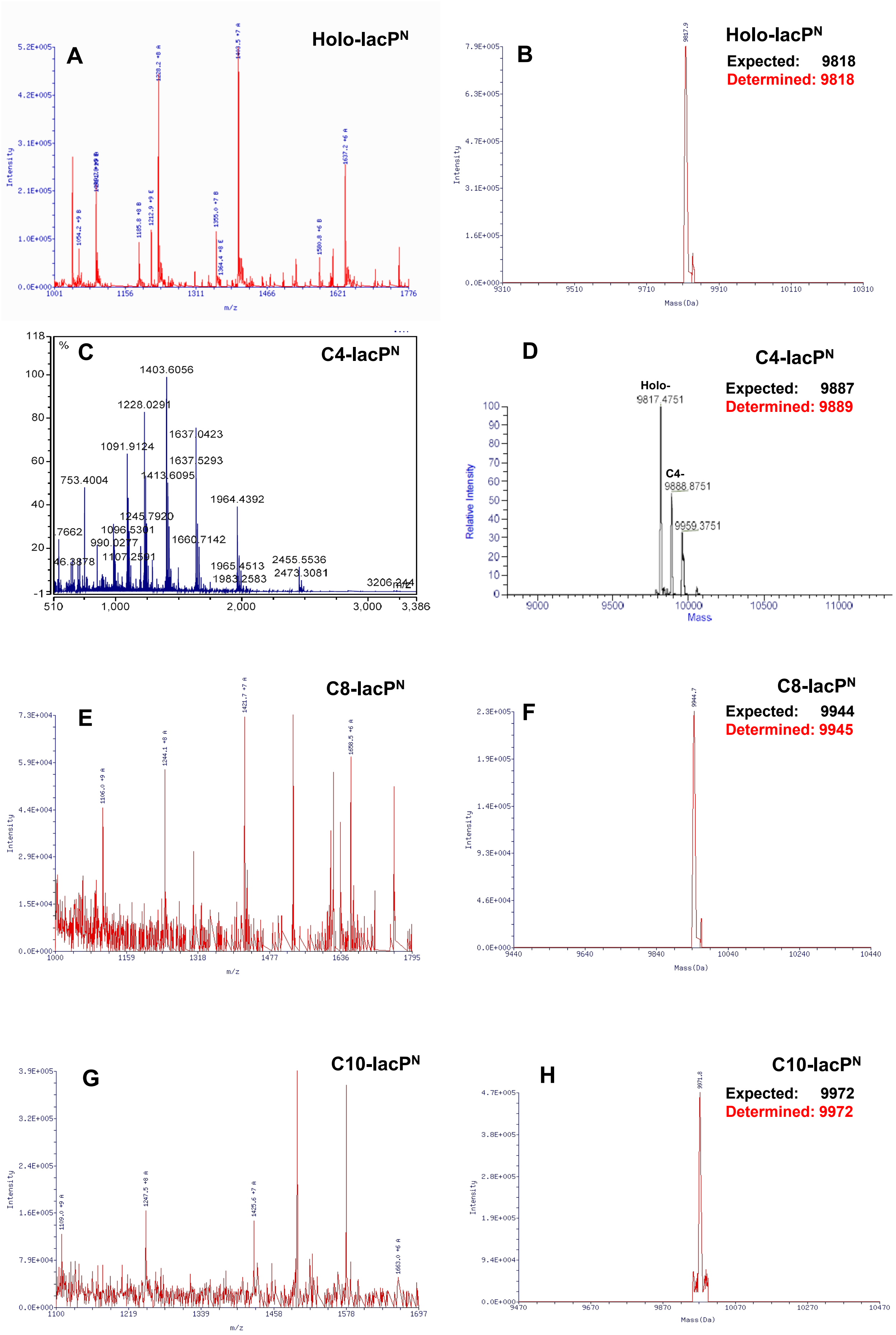
A
Holo-lacP^N
B
Holo-lacP^N
Expected: 9818
Determined: 9818
C
C4-lacP^N
1403.6056
1228.0291
1637.0423
1091.9124
1637.5293
753.4004
1413.6095
1964.4392
1245.7920
1096.5301
990.0277
1660.7142
1107.2591
1965.4513
2455.5536
1983.2583
2473.3081
D
C4-lacP^N
Holo-
9817.4751
Expected: 9887
Determined: 9889
C4-
9888.8751
9959.3751
Relative Intensity
Mass
E
C8-lacP^N
F
C8-lacP^N
Expected: 9944
Determined: 9945
G
C10-lacP^N
H
C10-lacP^N
Expected: 9972
Determined: 9972
Intensity
m/z
Mass(Da)

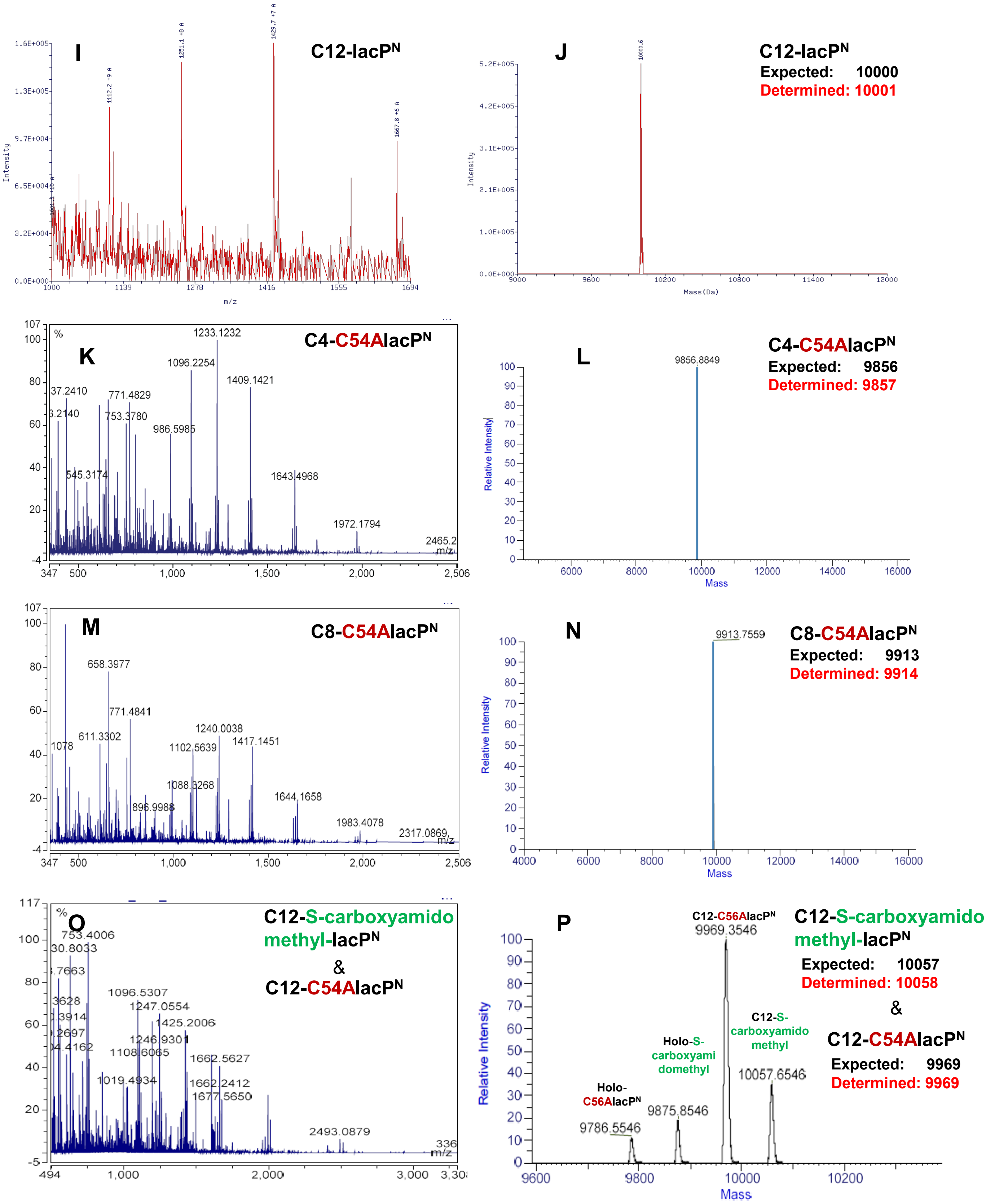


**Fig. S3 ESI-MS spectra of the acyl-lacP$^N$ intermediates.**
Representative spectra showing both raw data and the deconvoluted envelopes for A&B) holo-lacP$^N$, C&D) C4-lacP$^N$, E&F) C8-lacP$^N$, G&H) C10-lacP$^N$, I&J) C12-lacP$^N$ intermediates, K&L) C4-C54AlacP$^N$, M&N) C8-C54AlacP$^N$, O&P) a mixture of C12-C54AlacP$^N$ and C12-S-carboxyamidomethyl-lacP$^N$.

In the spectral region P), the peaks corresponding to the two intermediates C12-C56AIacP$^N$ and C12-S-carboxyamidomethyl-IacP$^N$ are labeled. In both cases, a fraction of each of the intermediates appeared hydrolyzed and therefore the envelope for the holo- peaks are also visible. From the mass it appears that only one cysteine is getting modified with IAA in C12-S-carboxyamidomethyl-IacP. The determined masses were within ±1 Da of the expected value. IacP is extremely hydrophobic, and all of the protein that binds to the C4 column does not elute. Therefore, the intensity of the signal is relatively low in most of the spectra (raw data).

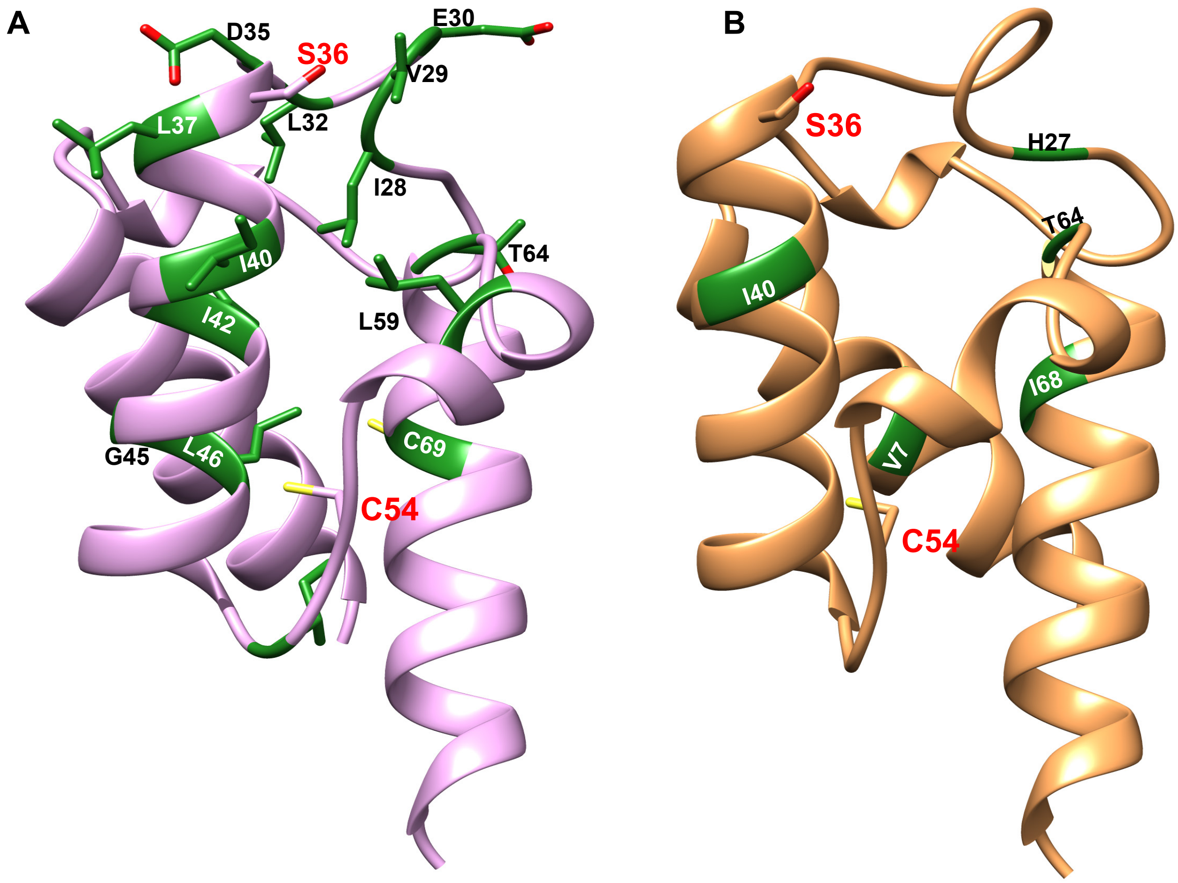


**Fig. S4 Residues displaying peak twinning in the acyl-lacP$^N$ intermediates are present in loop I-helix IV.**

A) The residues of lacP that display peak twinning in the acyl-lacP$^N$ intermediates, and B) residues that display noticeable C$\alpha$ change in the C8- and C10-acyl intermediates followed in an HNCA experiment.

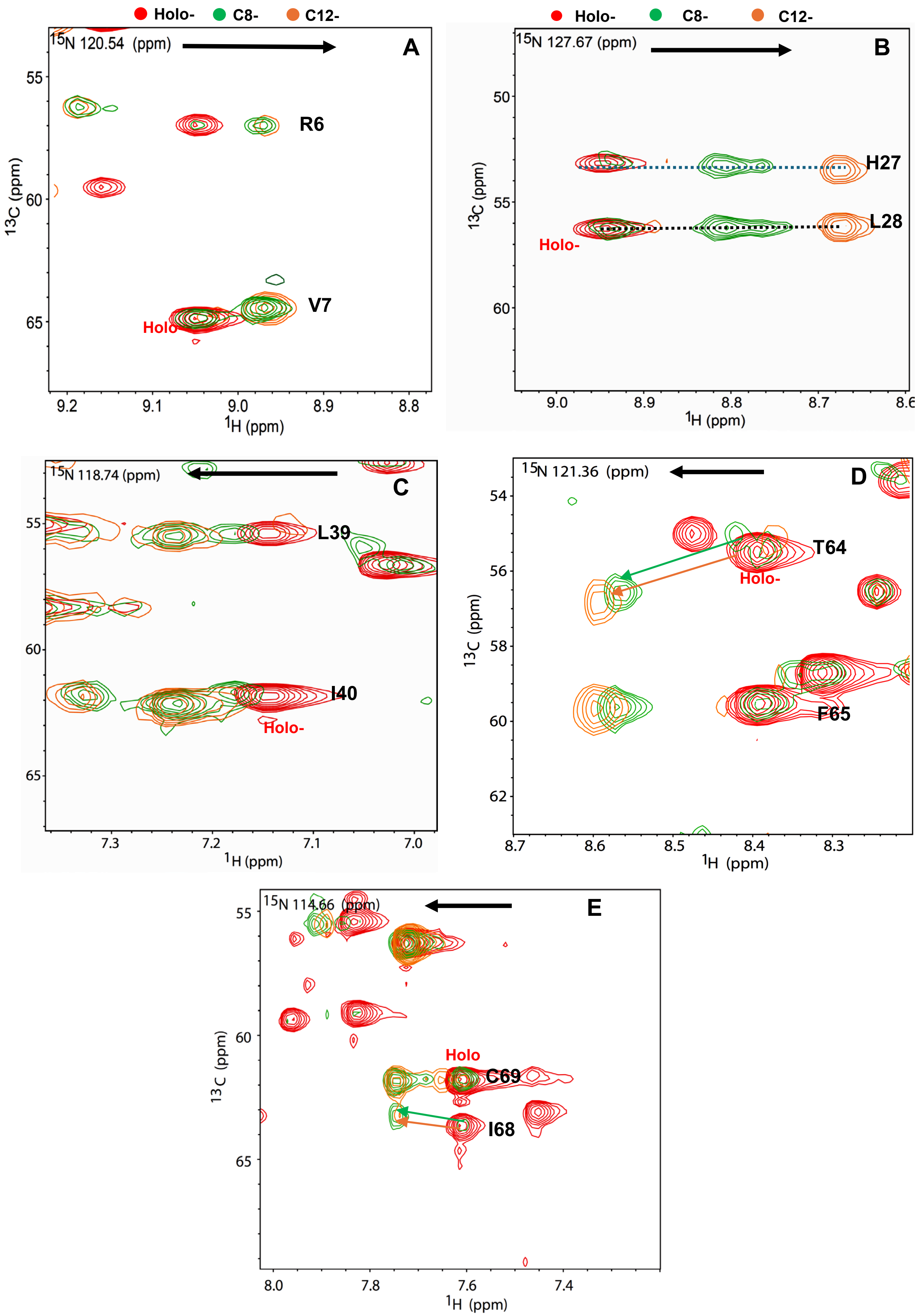


**Fig. S5 Significant Cα perturbations were observed for a few of the residues in the acyl-lacP$^{N}$ intermediates.** The $^{15}N^{13}C$ HNCA spectra for holo-lacP$^{N}$ (red), C8- (green), and C10- (brown) lacP$^{N}$ intermediates.
